# “Neutral and niche dynamics in a synthetic microbial community”

**DOI:** 10.1101/107896

**Authors:** NJ Cira, MT Pearce, SR Quake

**Affiliations:** Department of Bioengineering, Stanford University, Stanford, CA 94305.; Department of Physics, Stanford University, Stanford, CA 94305.

## Abstract

Ecologists debate the relative importance of niche versus neutral processes in understanding biodiversity^1,2^. This debate is especially pertinent to microbial communities, which play crucial roles in biogeochemical cycling^3,4^, food production^5^, industrial processes^6,7^, and human health and disease^8^. Here we created a synthetic microbial community using heritable genetic barcodes and tracked community composition over time across a range of experimental conditions. We show that a transition exists between the neutral and niche regimes, and, consistent with theory, the crossover point depends on factors including immigration, fitness, and population size. We find that diversity declined most rapidly at intermediate population sizes, which can be explained by a tradeoff between replacement by migration and duration of growth. We then ran an experiment where the community underwent abrupt or gradual changes in size, the outcome of which highlights that selecting the correct model is essential to managing diversity. Taken together these results emphasize the importance of including niche effects to obtain realistic models across a wide range of parameters, even in simple systems.

## Main text

Microorganisms are essential players in areas such as health, disease, industry, and the environment. Furthermore, it is often the community that gives rise to the output or property of interest^9^, rather than any individual organism. Understanding microbial communities is therefore important in a wide variety of systems for predicting responses to anthropogenic^4^ and natural perturbations, engineering desired outputs^10,11^, and understanding native functions. The study of microbial communities has been aided by increasing quantities of data as sequencing technologies have rapidly advanced, but in order to move from taxonomic descriptions to deeper understanding, there have been calls for placing these results in the context of a theoretical framework^12–18^.

Borrowed from macroscopic ecology, theoretical frameworks of microbial ecology can be divided into two types: models based on niche theory and neutral models. Models based on niche theory take a wide variety of forms, but critically differ from neutral theories in that they specify explicit differences between community members. For instance, one species may grow faster in certain abiotic conditions or might be killed as prey to feed another species^19^. These models can make detailed predictions, but often necessitate measurements or estimates of many parameters^20,21^. Conversely, neutral models take into account only random mechanisms. In ecology, they draw no distinction between individuals even across different species, which act in competition for a single limiting resource. Each species has the same fitness, and each species’ relative abundance changes only due to random processes such as immigration and drift from random sampling^1^. Despite obvious species differences documented by decades of observation, neutral models can, perhaps surprisingly, predict frequently observed patterns of natural communities, such as lognormal-like species relative abundance distributions^22^ and power law-like species area relationships^23^. Neutral models are thus a potentially enticing way to understand communities by abstracting away complicated differences between species, but a natural question arises about their applicability under varying balances between niche and random processes.

As in macroscopic ecology, both niche and neutral theories have been discussed in microbial ecology, and studies have reached a wide range of conclusions about the relative contributions of niche and neutral processes to community assembly and structure^24–35^. As a result, researchers have emphasized the need for controlled time course experiments to explore how the balance between different mechanisms such as selection, drift, and immigration compels the application of different models^14,36^. Toward this end, we created a synthetic microbial “community” to emulate an ecological community. In this community “species” are represented by unique heritable DNA barcodes^37,38^ that distinguish otherwise clonal *E. coli* bacterial cells. We created and validated a library of 456 different strains with Sanger sequencing to become the different species in our community. Starting with all species present, we grew this community to saturation then passaged it through a bottleneck once per day in 2 mL of shaken rich media using a wide range of bottleneck sizes (~10^0^ to 10^7^ cells, referred to as “small” and “large” bottlenecks, respectively). After each bottleneck we immigrated a controlled number of cells from the naïve barcoded “metacommunity” (average 55 cells/round). We took samples from the saturated growth at each time point, and used high throughput amplicon sequencing of the inserted barcodes to measure the abundance of each species present in each experimental community every round for 25 days. (Figure 1). Our approach allowed detection of species down to abundances of 1/1,000, creating a detection threshold analogous to Preston’s veil^39^. Details of the experimental methods can be found in the Supplementary methods section.

**Figure 1.**
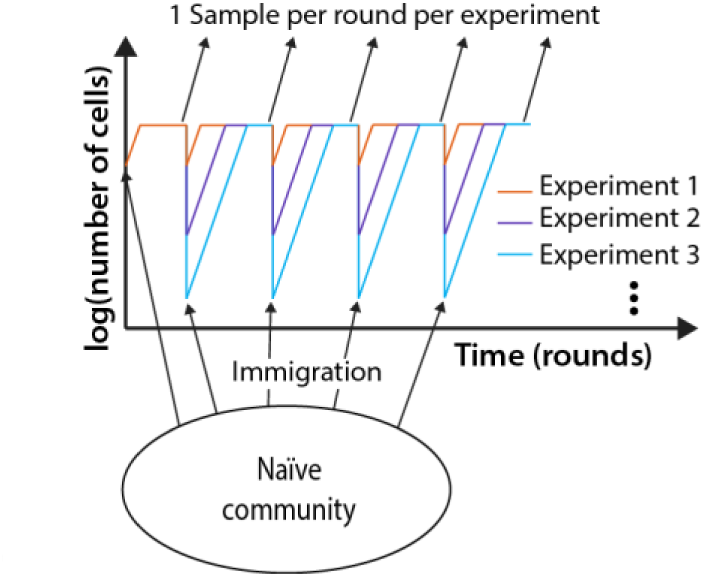
Experimental setup. 456 genetically barcoded *E. coli* strains were serially propagated at a variety of dilutions with an influx of immigrant cells after each dilution.

We note, of course, that this simplified system far from captures the complexity present in natural microbial communities. The species exist in a well-mixed environment and, since they started as clones, they share similar fitness, and the same nutrient requirements. Therefore, interactions between species are essentially a zero-sum competition for a limiting resource and do not include other types of interactions such as mutualism or predation. Recall, however, that neutral models only account for competitive interactions between species, and do not account for fitness differences. In fact, deviations from these conditions might be expected to drive the system away from neutral dynamics, so the experiments here might be expected to be even more neutral than higher complexity natural communities.

Figure 2A shows the number of species present over time in nine different experiments as the bottleneck size is varied. We find that the number of species present in each condition declines from the initial state with a variety of dynamics depending on bottleneck size. For each experiment we can also visualize the relative abundance of all 456 species over time with Muller plots (Figure 2 C-E, Supplementary figure 12) which show differences in dynamics between different species within a single experiment and different patterns of dynamics across bottleneck sizes. In analogy to classic species area curves^40,41^, where area is often assumed to be proportional to the number of individuals^41^, we also create log-log plots of the number of species present as a function of number of individuals passing through the bottleneck. These plots change with time and do not appear to reach an equilibrium (Figure 2B). We also visualize the data with residence time histograms, rank abundance plots, and species relative abundance histograms. (Supplementary figures 15, 18, 21). For a simple case with no immigration see Supplementary figure 8.

**Figure 2.**
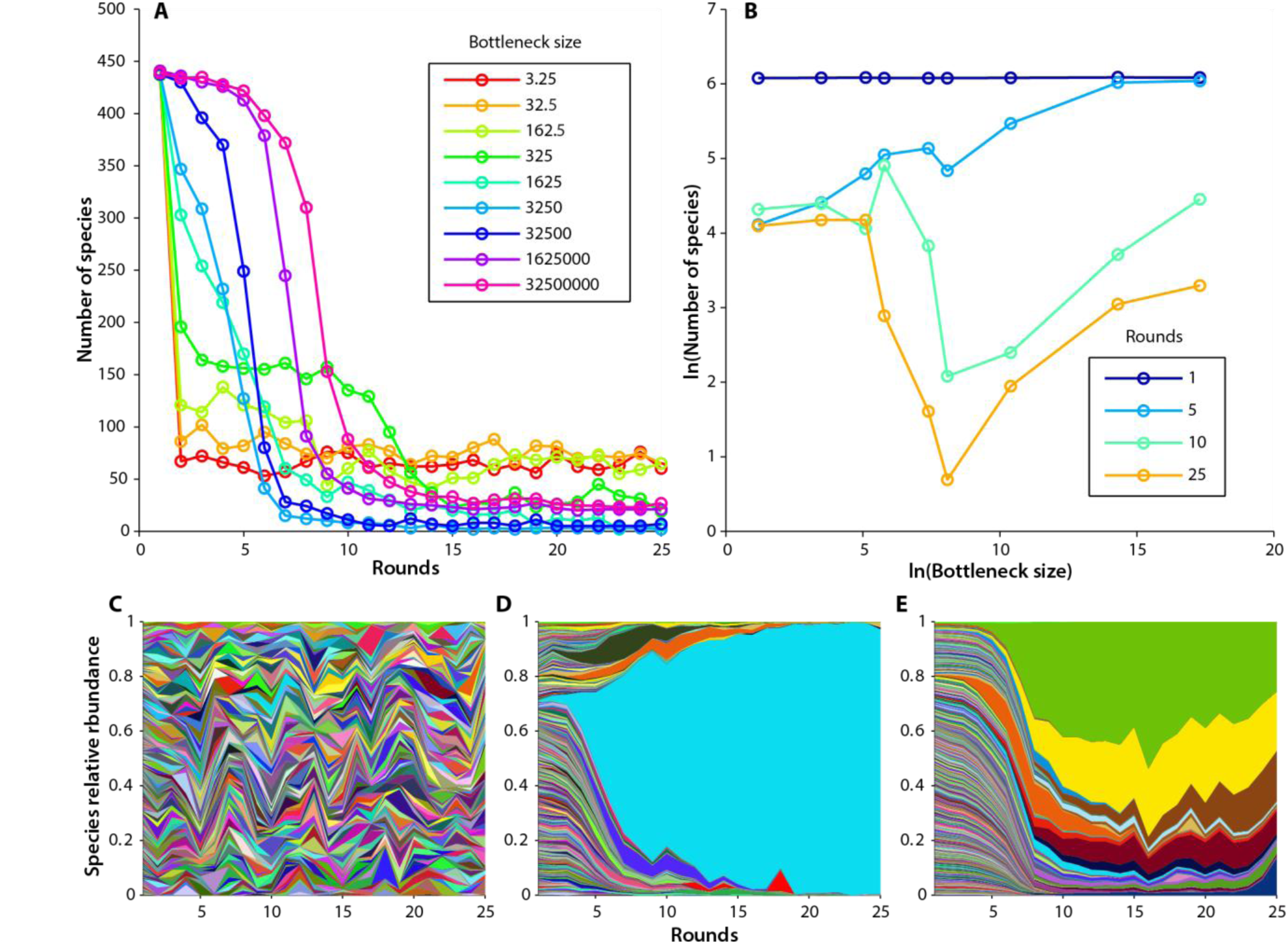
Experimental time dynamics. Immigration is ~55 cells/round for each experiment. A) Number of species detected over time for a range of dilutions. B) Species area curves over time. C-E) Muller plots showing the relative abundance of all 456 strains over time in three different experiments. Each species is represented by a different color where the proportion of each color represents the relative abundance of that species. Color/species pairings are consistent from plot to plot. C) bottleneck = 3.2 cells/round. D) bottleneck = 1625 cells/round. E) bottleneck = 1,625,000 cells/round.

We constructed and simulated a simple neutral model in an attempt to capture the system dynamics. The simulation has 25 rounds for each time point of the experiment. Each round, the new community is chosen from the old community by Poisson sampling to account for the bottleneck size, *N*. After the bottlenecking event, a mean number of cells, *M*, are immigrated from the original naïve population to the community, also by a Poisson process. When there is no immigration, this neutral model predicts that the community will eventually contain only one species (Supplementary figure 8). When immigration is included the number of species present eventually reaches a stable equilibrium (Figure 3A, Supplementary figure 10) independent of starting conditions (Supplementary figure 9). Like other neutral models^23^, this model predicts a power law relationship with exponent near ¼ when the number of species present is plotted against the bottleneck size for simulations that reach equilibrium (Figure 3I). Characteristic species relative abundance plots are also predicted. (Supplementary figure 22) Details of the model and expanded results can be found in Supplementary sections 2 and 3. A neutral model with additional stochasticity such as a very large variance in growth could lead to a fast loss of diversity as seen in the experiments, however our experimentally extracted estimates of the growth variance in individual lineages show that this variance is far too small to explain the loss of diversity at large bottleneck sizes (Supplementary section 4).

**Figure 3.**
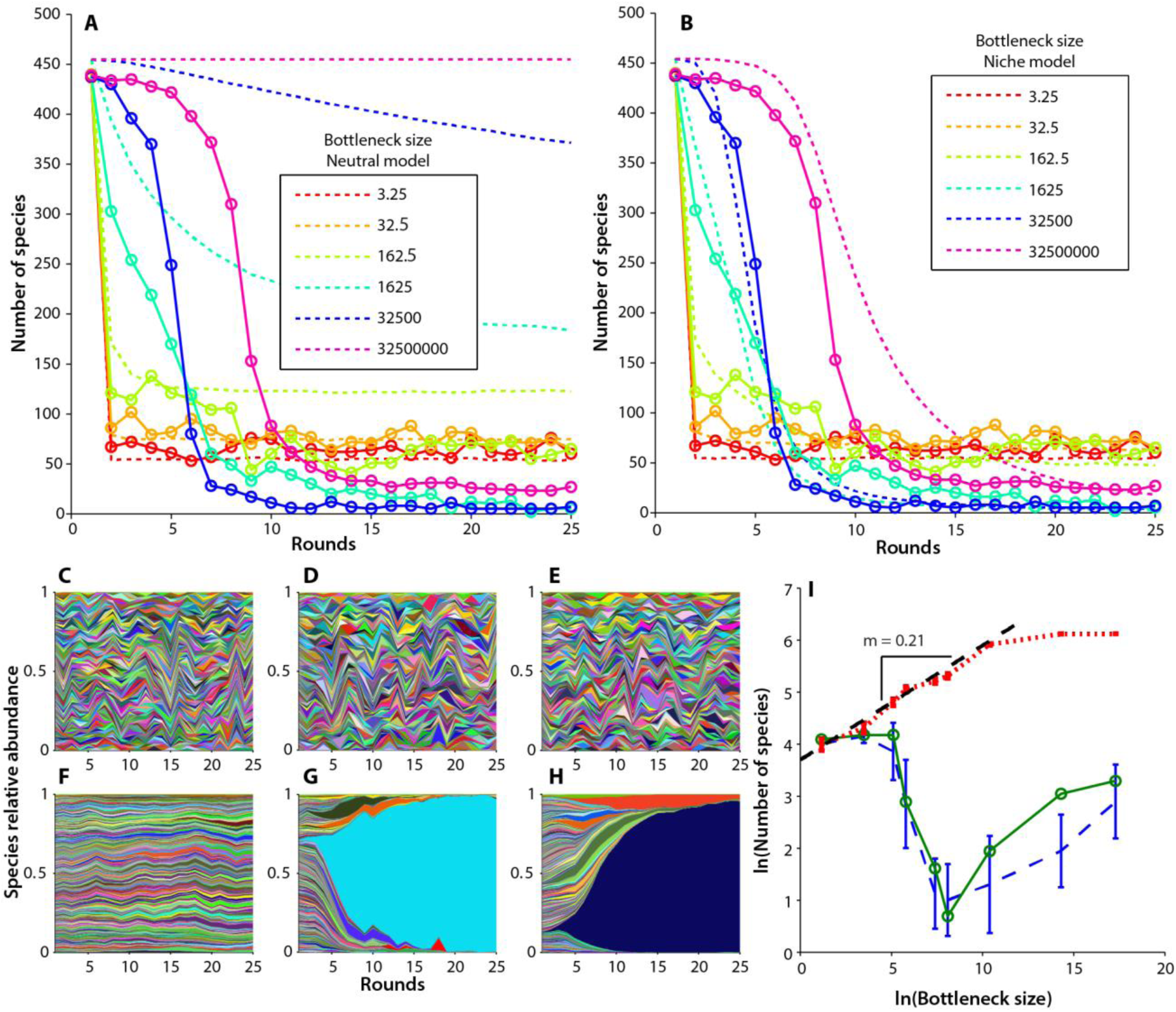
Comparison of neutral and niche models to experimental data. A-B) Number of species detected over time for a range of bottleneck sizes. Solid lines are experiment, dashed lines are model (means of 100 simulations). Note, for ease of visualization, not all bottleneck sizes are displayed. For the remaining bottleneck sizes, see Supplementary section 3 A) Neutral model. B) Niche model. C-H) Muller plots showing the relative abundance of all 456 strains over time for two different bottleneck sizes (representative trials picked for simulations) C-E) bottleneck = 3.25 cells/round F-H) bottleneck = 1625 cells/round for neutral (C & F) and niche (E & H) models compared to experiment (D & G). I) End point (round 25) species area curves for neutral (red dashed line), and niche (blue dashed line) models compared to experiment (solid green line). Note the neutral model predicts a power law relationship for the smaller bottleneck sizes. Error bars denote one standard deviation from 100 simulated trials.

Community dynamics in many experiments begin to differ drastically from the predictions of the neutral model (Figure 3A). During the early time points (rounds 1-5) experiments with larger bottlenecks lose diversity slower relative to those with smaller bottlenecks as predicted by the neutral model, but experiments with medium and large bottlenecks lose diversity at much faster absolute rates than predicted. At later time points, the experiments with the two smallest bottlenecks continue to match the neutral predictions well, but experiments with medium sized bottlenecks have the lowest diversity, and experiments with large bottlenecks have lost an intermediate amount of diversity. This results in an experimental species vs area plot which does not follow a monotonic trend or stabilize over the duration of the experiment and rapidly develops a pronounced minimum at medium bottleneck sizes contrasting sharply with the neutral prediction. (Figure 3I)

A comparison of the Muller plots between the experiment and the neutral model illuminates the cause of the discrepancy. Muller plots from the neutral model match the experiments with the smallest two bottlenecks reasonably well (Figure 3C vs 3D). However, in all experiments with larger bottlenecks the neutral model predicts relatively uniform and consistent relative abundances between species through the simulated time, compared to the experiments where one or more species begin to take over the population. (Figure 3F vs 3G, Supplementary figures 12&13) As these species become dominant, the community rapidly loses diversity. The dominant species seem to rise in prevalence exponentially over time (Figure 3G), typical of a fitness advantage instead of a random process. For species that appeared to grow adaptively, we extracted their relative fitnesses from the relative abundance vs time data, correcting for immigration. We obtained maximum per round Malthusian relative fitnesses, *R*, of 100-180%, translating to maximum per replication Malthusian relative fitnesses, *r*, of 5-15%, *R* = *xr*, where *x* is the number of replication cycles required to grow to *N*_*f*_, the final population size, given by 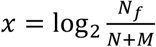. *R* decreased approximately linearly with log *N*, consistent with constant *r* across experiments due to an advantage in the exponential phase of growth. We noticed overlap in the identities of the fit species across experiments, suggesting the presence of preexisting fitness differences. Regardless of the underlying source of these fitness differences (For a discussion of these see Supplementary section 5, Supplementary table 1) they appear to cause deviations from the neutral predictions for larger bottleneck sizes.

A more complex model that captures the dynamics over a larger range of bottleneck sizes might then depart from neutrality and include fitness differences between species. We changed the model to include preexisting fitness differences by assigning each species a per replication relative fitness, constant across experiments, selected randomly from an exponential distribution^42,43^. We then scaled this fitness by the number of replication rounds per experiment to obtain a per round fitness for each species, consistent with advantages in the exponential growth phase (Supplementary section 5). This modification captured many more features over a larger range of the parameter space (Figure 3B) including species relative abundance trajectories where one or more species come to dominate (Figure 3H) and non-monotonic species area curves which do not stabilize over the experiment (Figure 3I). Further additions to the model such as including the chance for mutations to arise during the course of the experiment may lead to a more complete picture, especially at timescales beyond those investigated here. Noting the success of the neutral model at small bottlenecks, we next assess when additional complexities departing from neutrality become necessary.

The fact that small fitness differences can lead to non-neutral dynamics has been understood in the population genetics literature^44^ for some time and has more recently been studied in the context of neutral ecology models^24,45–47^. Transitions between niche and neutral have been proposed along speciation^48^ and immigration^24^ gradients and with different interplays between species interactions and stochasticity^36,49–51^. In our experiment, the different bottleneck sizes have different proportions of immigrants, 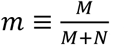, allowing us to explore the transition from neutral to niche along an immigration gradient in a well-controlled experiment. We can compare our results to theoretical predictions for the simple scenario of a single species with a fitness advantage, *R*. Our discussion of the predictions follows the derivation by Sloan *et al.*^24^ and can be found in Supplementary section 2c.

For the neutral case, any given species’ mean frequency, is simply equal to that species’ frequency in the incoming immigrant pool, *f*_*m*_. When there is a selective advantage, the probability distribution of the fit species, *p*(*f*), is shifted toward higher frequencies. This effect is most noticeable when selection is stronger than stochastic effects, i.e 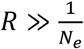 where *N*_*e*_ is the effective population size which is on the order of the total population after immigration, *N* + *M*. Even if selection is strong, the distribution can appear neutral if immigration is strong enough to compensate. For strong selection and strong immigration, the equilibrium frequency, *f*_*eq*_, of the fit species given by the deterministic dynamics is a good measure for determining whether the distribution appears neutral or not. Figure 4A shows *f*_*eq*_ as a function of the selective advantage, *R*, and immigration proportion, *m*, with the experimentally investigated points noted. The equilibrium frequency transitions from *f*_*m*_ when neutral up to 1 (indicating near-fixation) when non-neutral.

**Figure 4.**
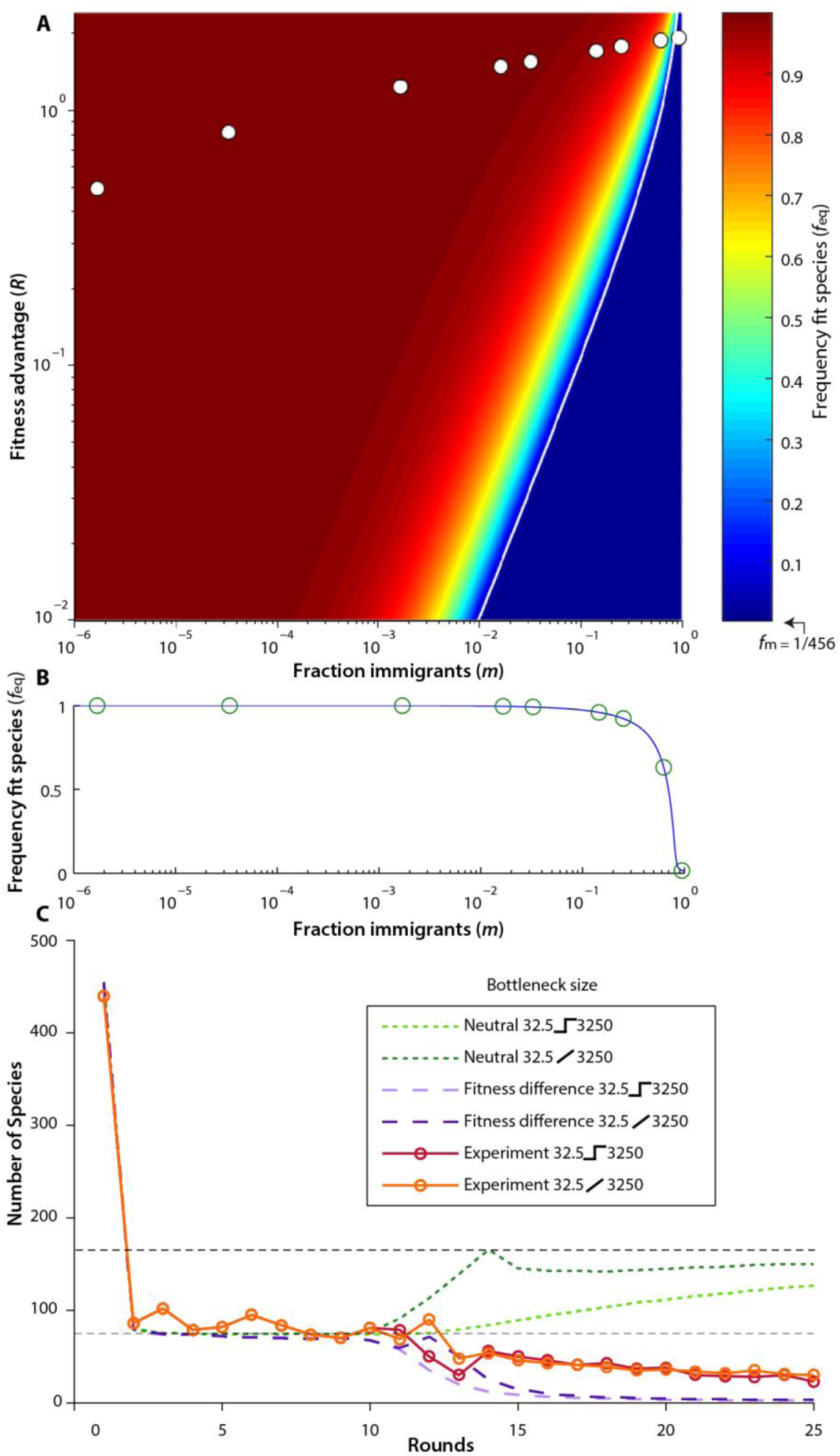
Transition from niche to neutral. A) Phase space of equilibrium frequency (*f*_*eq*_) as a function of per round relative fitness (*R*) vs fraction of immigrants (*m*). Points indicate the experimentally investigated region where bottleneck size decreases from left to right (assuming a maximum per replication relative fitness of 8.5%). The immigration fraction of any species is, *f*_*m*_ ≈ 1/456. The white line indicates the *R* = *δ* threshold. B) A slice through the phase space along the experimentally tested conditions. A transition is predicted: at high immigrant fraction fit clones do not rise to high abundance and the system is neutral, and at the low immigration fraction the fit species dominates the population and cause departure from neutrality. Each circle indicates the theoretical prediction for *f*_*eq*_. C) Community recovery. Here a community is maintained at a bottleneck size of 32.5 for 10 rounds then the bottleneck is allowed to expand to 3250. The recovery took the form of either a step function, or a gradual expansion. Though both models predict a similar number of species to the experimental community before recovery (lower horizontal dashed line), the neutral model (green lines) makes drastically different predictions than a model incorporating fitness differences (purple lines) after recovery. The neutral model predicts that the number of species in the community will increase to the new equilibrium level (upper horizontal dashed line), with the recovery happening much slower for the step function (light green dashed line) than the gradual increase (dark green dashed line). The model incorporating fitness differences predicts the community will lose diversity independent of step (lavender dashed line) or gradual recovery (purple dashed line). In both the step (solid red line) and gradual recovery (solid orange line) the experimental communities lost diversity.

The transition happens when selection and immigration are the same scale. It can be understood heuristically by considering the effective growth rate of cells already in the population. Immigration exerts an effective negative fitness effect because cells are replaced by new immigrants. The frequency would drop to 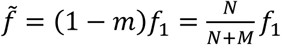 because the fraction *m* are replaced. It is convenient to define a negative fitness *δ* such that 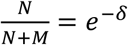. After immigration, the population grows again until the end of the cycle. The fit species’ frequency is then 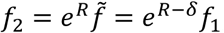, so *R* − *δ* acts as an effective fitness. If *R* − *δ* ≳ 0 then the frequency of cells in the population increases despite replacement by immigration. If we assume that the fitness advantage is an increase in the bacterial doubling rate by *r*, such that the fitness per cycle, *R*, scales with the number of doublings, then we can predict when the transition occurs for our experiments by solving *R* = *δ*. For *r* ≈ 8.5% (and using *M* = 55 and a final cell count of 6.5 × 10^9^) The transition is predicted when *m* ≈ 0.89. This compares well with the results that the smallest bottleneck (*m* ≈ 0.93) appears neutral while the second smallest bottleneck (*m* ≈ 0.63) and all larger bottlenecks appear non-neutral.

An interesting feature of the experiments is that the fastest exponential takeover and loss of diversity in the population happens at an intermediate bottleneck size, leading to, at least transiently, non-monotonic species area curves. If bottlenecking events are understood as a disturbance our results over this range of bottleneck sizes stand in contrast to the intermediate disturbance hypothesis, which suggests that diversity is maximized at intermediate levels of disturbance^52,53^. The concept of an effective fitness is useful in explaining this feature of the data; as the bottleneck size increases, the chance of being replaced by an immigrant (negative fitness effect) decreases, but the growth phase advantage (positive fitness effect) also decreases. This results in a tradeoff where the effective fitness is maximized when 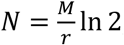, as found in Supplementary section 2c. With the same *M* and *r* as before, this gives *N* = 450, in agreement with the observation that a species in the *N* = 325 bottleneck had the largest effective fitness extracted from the experiments.

In our final experiment we addressed whether knowing which model to apply has practical implications for understanding and managing how the community recovers from a disturbance. Communities that have been restricted by severe bottlenecks lose diversity, for example the human gut microbiome after antibiotic treatment^54^. A neutral model can make predictions about the best way to recover diversity. In certain parameter regimes, diversity is actually predicted to recover faster from a slow, rather than abrupt, increase in the bottleneck size. For example, in our system, neutral models predict that a community maintained with a bottleneck size of 32 recovering to a bottleneck size of 3250 with immigration of 55 individuals per round gains diversity faster when the bottleneck is increased over several rounds rather than in a single round, prescribing a management strategy where the population gradually expands in size to maximize the rate at which diversity is recovered. (Figure 4C) Drastically different from neutral predictions, a model including fitness differences between species predicts that diversity would only continue to decay as the bottleneck size was increased, regardless of the dynamics of the increase, since increasing the bottleneck size decreases the immigration fraction, thereby increasing the effective fitness and only helping the fit species to outcompete. The experiment reveals that as the bottleneck is relaxed, regardless of relaxation dynamics, diversity is lost, in far better agreement with the model incorporating fitness differences, since the system transitioned well out of the regime where a neutral model was appropriate.

These results show a transition between niche and neutral regimes, providing an experimental case where a general guideline using fitness differences and immigration proportion successfully predicts whether the system can be treated as neutral. If these conditions are not met, then non-neutral explanations are required to understand the community. These results also show that using the correct model is essential when predicting community response to change and can impact management strategies. Finally, we note that though these results were obtained using a synthetic microbial community, the framework, models, and analytical results may be useful in other ecological systems involving fitness differences and immigration.

## Acknowledgements

We gratefully acknowledge S. Norviel for assistance in validating the barcoded library, DS Fisher, JR Blundell, A Agarwala, F. Zanini, GR Dick, J Russel, and JT Morton for helpful comments, and C Vollmers, GR Mantalas, NF Neff, J Okamoto, and B Passarelli for help with sequencing and informatics. NJC and MTP are supported by NSF GRFP Fellowships, NJC is supported by a Siebel Scholar Fellowship, and MTP is supported by a Sara Hart Kimball Fellowship as part of Stanford’s Graduate Fellowship Program. This research was supported by NASA Exobiology grant EXO-NNX11AR78G and U.S. National Science Foundation grants MCB 0546865, OISE 0968421, and DGE-114747.

## References

1. Hubbell, S. P. The Unified Neutral Theory of Biodiversity and Biogeography. 17, (Princeton University Press, 2001).

2. Chave, J., Muller-Landau, H. C. & Levin, S. A. Comparing classical community models: theoretical consequences for patterns of diversity. Am. Nat. 159, 1–23 (2002).

3. Francis, C. A., Beman, J. M. & Kuypers, M. M. New processes and players in the nitrogen cycle: the microbial ecology of anaerobic and archaeal ammonia oxidation. ISME J 1, 19–27 (2007).

4. Bardgett, R. D., Freeman, C. & Ostle, N. J. Microbial contributions to climate change through carbon cycle feedbacks. ISME J 2, 805–814 (2008).

5. Ardhana, M. M. & Fleet, G. H. The microbial ecology of cocoa bean fermentations in Indonesia. Int. J. Food Microbiol. 86, 87–99 (2003).

6. Wagner, M. et al. Microbial community composition and function in wastewater treatment plants. Antonie Van Leeuwenhoek 81, 665–680 (2002).

7. Saunders, A. M., Albertsen, M., Vollertsen, J. & Nielsen, P. H. The activated sludge ecosystem contains a core community of abundant organisms. ISME J. 10, 11–20 (2016).

8. Tremaroli, V. & Bäckhed, F. Functional interactions between the gut microbiota and host metabolism. Nature 489, 242–249 (2012).

9. Lepp, P. W. et al. Methanogenic Archaea and human periodontal disease. Proc. Natl. Acad. Sci. U. S. A. 101, 6176–81 (2004).

10. Shou, W., Ram, S. & Vilar, J. M. G. Synthetic cooperation in engineered yeast populations. Proc. Natl. Acad. Sci. U. S. A. 104, 1877–82 (2007).

11. Fredrickson, J. K. Ecological communities by design. Science (80-. ). 348, 1425–1427 (2015).

12. Prosser, J. I. et al. The role of ecological theory in microbial ecology. Nat. Rev. Microbiol. 5, 384–392 (2007).

13. Faust, K. & Raes, J. Microbial interactions: from networks to models. Nat. Rev. Microbiol. 10, 538–550 (2012).

14. Hanson, C. A., Fuhrman, J. A., Horner-Devine, M. C. & Martiny, J. B. H. Beyond biogeographic patterns: processes shaping the microbial landscape. Nat. Rev. Microbiol. 10, 1–10 (2012).

15. Robinson, C. J., Bohannan, B. J. M. & Young, V. B. From structure to function: the ecology of host-associated microbial communities. Microbiol. Mol. Biol. Rev. 74, 453–76 (2010).

16. Widder, S., Allen, R., Pfeiffer, T., Curtis, T. & Wiuf, C. Challenges in microbial ecology: building predictive understanding of community function and dynamics. ISME 10, 2557–2568 (2016).

17. Dethlefsen, L., McFall-Ngai, M. & Relman, D. A. An ecological and evolutionary perspective on humang-microbe mutualism and disease. Nature 449, 811–818 (2007).

18. Lennon, J. T. & Locey, K. J. Macroecology for microbiology. Environ. Microbiol. Rep. (2017). doi:10.1111/1758-2229.12512

19. Alfred J. Lotka. Elements of Physical Biology. (Williams and Wilkins Company, 1925). doi:10.2105/AJPH.15.9.812-b

20. Follows, M. J., Dutkiewicz, S., Grant, S. & Chisholm, S. W. Emergent biography of microbial communities in a model ocean. Science (80-. ). 315, 1843–1846 (2007).

21. Allison, S. D. A trait-based approach for modelling microbial litter decomposition. Ecology Letters 15, 1058–1070 (2012).

22. Hubbell, S. P. Tree dispersion, abundance, and diversity in a tropical dry forest. Science (80-. ). 203, 1299–1309 (1979).

23. Bell, G. The distribution of abundance in neutral communities. Am. Nat. 155, 606–617 (2000).

24. Sloan, W. T. et al. Quantifying the roles of immigration and chance in shaping prokaryote community structure. Environ. Microbiol. 8, 732–740 (2006).

25. Woodcock, S. et al. Neutral assembly of bacterial communities. FEMS Microbiol. Ecol. 62, 171–180 (2007).

26. Horner-Devine, M., Silver, J. & Leibold, M. A Comparison of taxon co‐occurrence patterns for macro‐and microorganisms. Ecology 88, 1345–1353 (2007).

27. Ayarza, J. M. & Erijman, L. Balance of Neutral and Deterministic Components in the Dynamics of Activated Sludge Floc Assembly. Microb. Ecol. 61, 486–495 (2011).

28. Langenheder, S. & Székely, A. Species sorting and neutral processes are both important during the initial assembly of bacterial communities. ISME J. 5, 1086–1094 (2011).

29. Zhang, Q. G., Buckling, A. & Godfray, H. C. J. Quantifying the relative importance of niches and neutrality for coexistence in a model microbial system. Funct. Ecol. 23, 1139–1147 (2009).

30. Ofiteru, I. D. et al. Combined niche and neutral effects in a microbial wastewater treatment community. Proc. Natl. Acad. Sci. U. S. A. 107, 15345–15350 (2010).

31. Jeraldo, P. et al. Quantification of the relative roles of niche and neutral processes in structuring gastrointestinal microbiomes. Proc. Natl. Acad. Sci. U. S. A. 109, 9692–8 (2012).

32. Dumbrell, A. J., Nelson, M., Helgason, T., Dytham, C. & Fitter, A. H. Relative roles of niche and neutral processes in structuring a soil microbial community. ISME J. 4, 337–345 (2010).

33. Fukami, T., Beaumont, H. J. E., Zhang, X.-X. & Rainey, P. B. Immigration history controls diversification in experimental adaptive radiation. Nature 446, 436–9 (2007).

34. Manefield, M., Whiteley, A., Curtis, T. & Watanabe, K. Influence of sustainability and immigration in assembling bacterial populations of known size and function. Microb. Ecol. 53, 348–354 (2007).

35. Venkataraman, A. et al. Application of a neutral community model to assess structuring of the human lung microbiome. MBio 6, (2015).

36. Fisher, C. & Mehta, P. The transition between the niche and neutral regimes in ecology. Proc. Natl. Acad. Sci. U. S. A. 111, 13111–13116 (2014).

37. Winzeler, E. A. et al. Functional characterization of the S. cerevisiae genome by gene deletion and parallel analysis. Science (80-. ). 285, 901–906 (1999).

38. Levy, S. F. et al. Quantitative evolutionary dynamics using high-resolution lineage tracking. Nature 519, 181–6 (2015).

39. Preston, F. W. The Commonness, And Rarity, of Species. Ecology 29, 254–283 (1948).

40. MacArthur, R. H. & Wilson, E. O. The theory of island biogeography. 1, (Princeton University Press, 1967).

41. Preston, F. W. The canonical distibution of commonness and rarity: Part I. Ecology 43, 182–215 (1962).

42. Orr, H. A. The Distribution of Fitness Effects Among Beneficial Mutations. Genetics 163, 1519–1526 (2003).

43. Kassen, R. & Bataillon, T. Distribution of fitness effects among beneficial mutations before selection in experimental populations of bacteria. Nat. Genet. 38, 484–8 (2006).

44. Fisher, R. A. The distribution of gene ratios for rare mutations. Proc. R. Soc. Edinb (1930).

45. Zhang, D. & Lin, K. The effects of competitive asymmetry on the rate of competitive displacement: how robust is Hubbell’s community drift model? J. Theor. Biol. 188, 361–367 (1997).

46. Yu, D., Terborgh, J. & Potts, M. D. Can high tree species richness be explained by Hubbell’s null model? Ecol. Lett. 1, 193–199 (1998).

47. He, F., Zhang, D. Y. & Lin, K. Coexistence of nearly neutral species. J. Plant Ecol. 5, 72–81 (2012).

48. Chisholm, R. A. & Pacala, S. W. Theory predicts a rapid transition from niche-structured to neutral biodiversity patterns across a speciation-rate gradient. Theor. Ecol. 4, 195–200 (2011).

49. Haegeman, B. & Loreau, M. A mathematical synthesis of niche and neutral theories in community ecology. J. Theor. Biol. 269, 150–165 (2011).

50. Pigolotti, S. & Cencini, M. Species abundances and lifetimes: From neutral to niche-stabilized communities. J. Theor. Biol. 338, 1–8 (2013).

51. Gravel, D., Canham, C. D., Beaudet, M. & Messier, C. Reconciling niche and neutrality: The continuum hypothesis. Ecol. Lett. 9, 399–409 (2006).

52. Connell, J. H. Diversity in Tropical Rain Forests and Coral Reefs. Science (80-. ). 199, 1302–1310 (1978).

53. Lubchenco, J. Plant Species Diversity in a Marine Intertidal Community: Importance of Herbivore Food Preference and Algal Competitive Abilities. Am. Nat. 112, 23–39 (1978).

54. Dethlefsen, L., Huse, S., Sogin, M. L. & Relman, D. A. The pervasive effects of an antibiotic on the human gut microbiota, as revealed by deep 16s rRNA sequencing. PLoS Biol. 6, 2383–2400 (2008).

